# Erythropoietin regulates transcription and YY1 dynamics in a pre-established chromatin architecture

**DOI:** 10.1101/2020.04.21.034116

**Authors:** Andrea A. Perreault, Jonathan D. Brown, Bryan J. Venters

## Abstract

The three dimensional architecture of the genome plays an essential role in establishing and maintaining cell identity. By contrast, the magnitude and temporal kinetics of changes in chromatin structure that arise during cell differentiation remain poorly understood. Here, we leverage a murine model of erythropoiesis to study the relationship between chromatin conformation, the epigenome, and transcription in erythroid cells. We discover that acute transcriptional responses induced by erythropoietin (EPO), the hormone necessary for erythroid differentiation, occur within an invariant chromatin topology. Within this pre-established landscape, Yin Yang 1 (YY1) occupancy dynamically redistributes to sites in proximity to EPO-regulated genes. Using HiChIP, we identify chromatin contacts mediated by H3K27ac and YY1 that are enriched for enhancer-promoter interactions of EPO-responsive genes. Taken together, these data are consistent with an emerging model that rapid, signal-dependent transcription occurs in the context of a pre-established chromatin architecture.

## INTRODUCTION

Transcription control is a primary mechanism for regulating gene expression in eukaryotes. Three major steps exist in the transcription cycle: 1) preinitiation complex (PIC) formation; 2) pause release of RNA Polymerase II (Pol II) to productive elongation; and 3) transcription termination(Liu et al., 2015). Multiple mechanisms exist to regulate each step, thereby providing precise control over the magnitude and kinetics of transcription and global gene expression. Promoter proximal pausing is one such mechanism and is recognized as a general feature of transcription at many eukaryotic genes. Specifically, there is a prominence of paused Pol II at signal responsive genes, which serves to prime these genes for rapid transcription in response to environmental stimuli(Adelman et al., 2009; Danko et al., 2013; Gaertner et al., 2012). TF-bound enhancers activate Pol II, acting as an additional mechanism in regulating transcription(Heintzman et al., 2009) and defining cell identity(Ernst et al., 2011; Zhu et al., 2013). While chromatin state maps are useful to assign enhancers to target genes based on distance from promoters, proximity analysis is overly simplistic with respect to the true gene-regulatory environment(Mora et al., 2016; Rao et al., 2014; Yao et al., 2015).

More recently, high-resolution maps of the three-dimensional (3D) genome have revealed that enhancers exhibit long-range control of transcription. Structural proteins, such as CCCTC-binding factor (CTCF) and Yin-Yang 1 (YY1), tether distal TF-bound enhancers to their target gene promoters. CTCF is an evolutionarily conserved zinc-finger that co-localizes with cohesin(Phillips and Corces, 2009). Together, these two factors establish and maintain chromatin loops(Rao et al., 2017; Ren et al., 2017; Sanyal et al., 2012). Assays that can map chromatin contacts, such as HiC, have revealed that the genome is organized into active and inactive domains, which are demarcated by CTCF(Dixon et al., 2012; Lieberman-Aiden et al., 2009) and cell type specific(Arzate-Mejia et al., 2018; Ong and Corces, 2014; Phillips and Corces, 2009). These large domains can be further separated into topologically associated domains (TADs) and subTADs that contain higher contact frequencies between regions of the genome, many of which are not limited to one-to-one interactions(Dowen et al., 2014; Fullwood et al., 2009; Ji et al., 2016; Li et al., 2012; Sanyal et al., 2012). Together, these findings demonstrate CTCF’s function as structural foci for chromatin organization, whereby Pol II can selectively target cell-type specific genes for transcription through interactions with looping factors and enhancers.

Other TFs, such as YY1, are specifically enriched at chromatin loops that connect enhancers to promoters of actively transcribed genes(Weintraub et al., 2017). YY1 is a ubiquitously expressed zinc-finger TF that plays an important role in cellular differentiation(Beagan et al., 2017; Kleiman et al., 2016). Deletion of YY1 binding motifs at gene promoters in mouse embryonic stem cells (ESCs) reduced contact frequency between individual promoters and enhancers, and variably reduced mRNA levels(Weintraub et al., 2017). These data provide evidence for an essential role of YY1 in controlling gene expression by facilitating E-P interactions.

Erythropoiesis has been a useful model system for understanding the interplay between Pol II dynamics(Johnson et al., 2002; Sawado et al., 2003), enhancer activity(Reik et al., 1998), and 3D genome structure(Bartman et al., 2016; Chien et al., 2011; Deng et al., 2012; Tolhuis et al., 2002) during cellular differentiation. Indeed, we have previously characterized the genome-wide enhancer landscape in proerythroblasts (ProEBs) in response to erythropoietin (EPO)(Perreault et al., 2017), the hormone that is required for terminal erythroid differentiation(Koury and Bondurant, 1988, 1990). However, the manner by which EPO signaling shapes the 3D genome and specific chromatin interactions remains poorly understood. Additionally, although CTCF occupancy and function has been assessed in erythrocytes(Hanssen et al., 2017; Hsu et al., 2017; Lee et al., 2017), the YY1 binding locations in erythroid cells are not known, resulting in a knowledge gap in uncovering the role of important TFs controlling E-P interactions and overall chromatin architecture during erythropoiesis.

To address this critical gap in understanding, we leveraged a murine model system to study synchronous erythroid maturation *ex vivo* in response to EPO stimulation (Fig. 1)(Bondurant et al., 1985; Koury et al., 1984; Sawyer et al., 1987). Here, we demonstrate that EPO stimulates rapid transcriptional changes in ProEBs after 1 hour (Fig. 2). During this time, YY1 occupancy is dynamically redistributed, as opposed to CTCF, which remains unchanged (Fig. 3). Moreover, there is little overlap in the regions bound by these structural TFs. Using HiChIP, we determined the chromatin contacts mediated by H3K27ac and YY1 genome-wide. We discover that a subset of these chromatin interactions remain invariant during EPO signaling, facilitating unique E-P interactions during EPO-mediated transcriptional regulation (Fig. 4).

**Figure 1.**
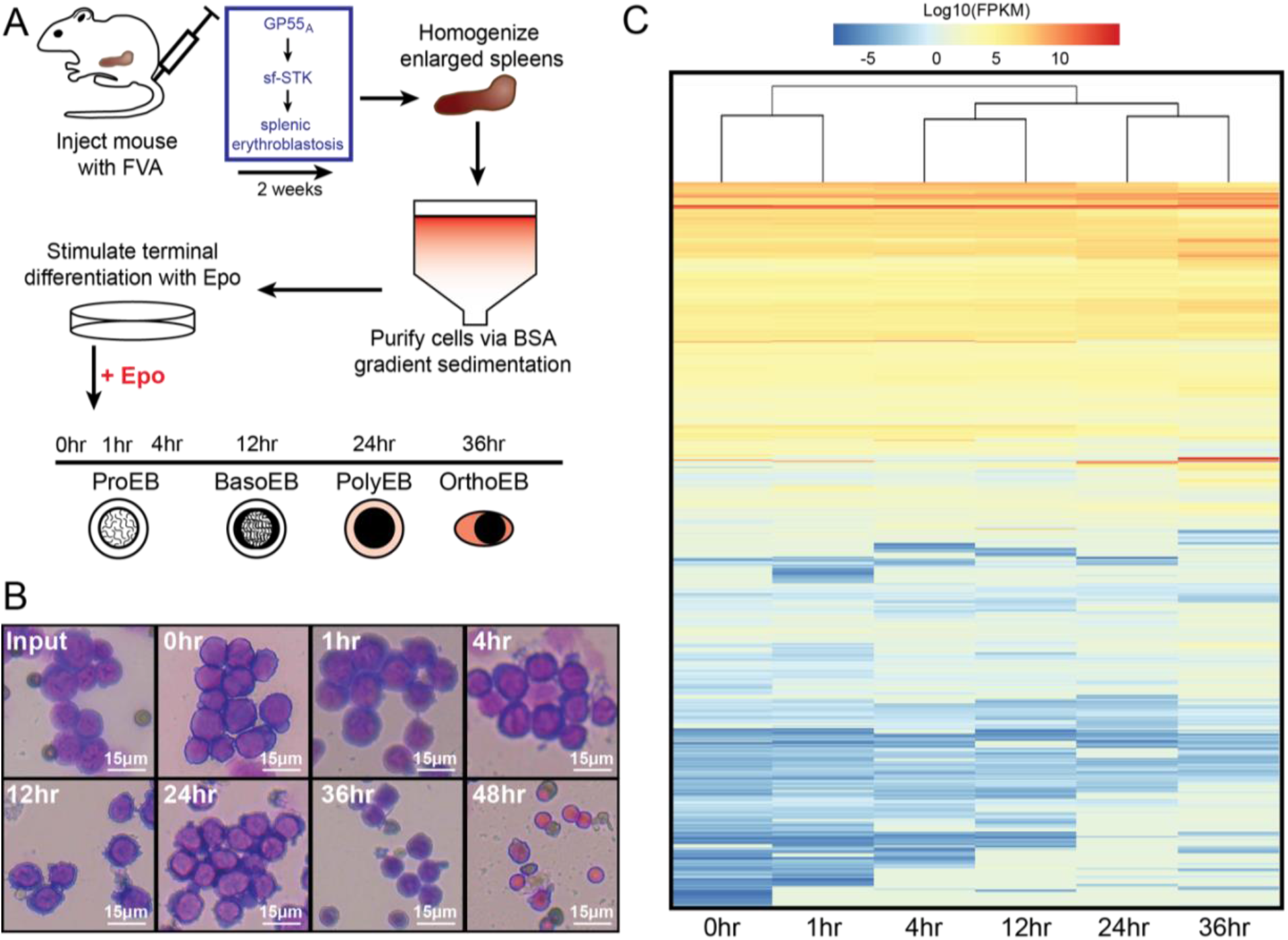
The FVA murine system faithfully recapitulates erythroid differentiation during erythropoiesis. (A) The workflow for generating and isolating highly purified EPO-responsive ProEBs from a mouse injected with the Friend Virus that induces Anemia (FVA). (B) Microscopy images highlighting morphological changes of ProEBs isolated using the FVA system during differentiation. (C) Heatmap of RNA-seq gene expression through erythroid differentiation.

**Figure 2.**
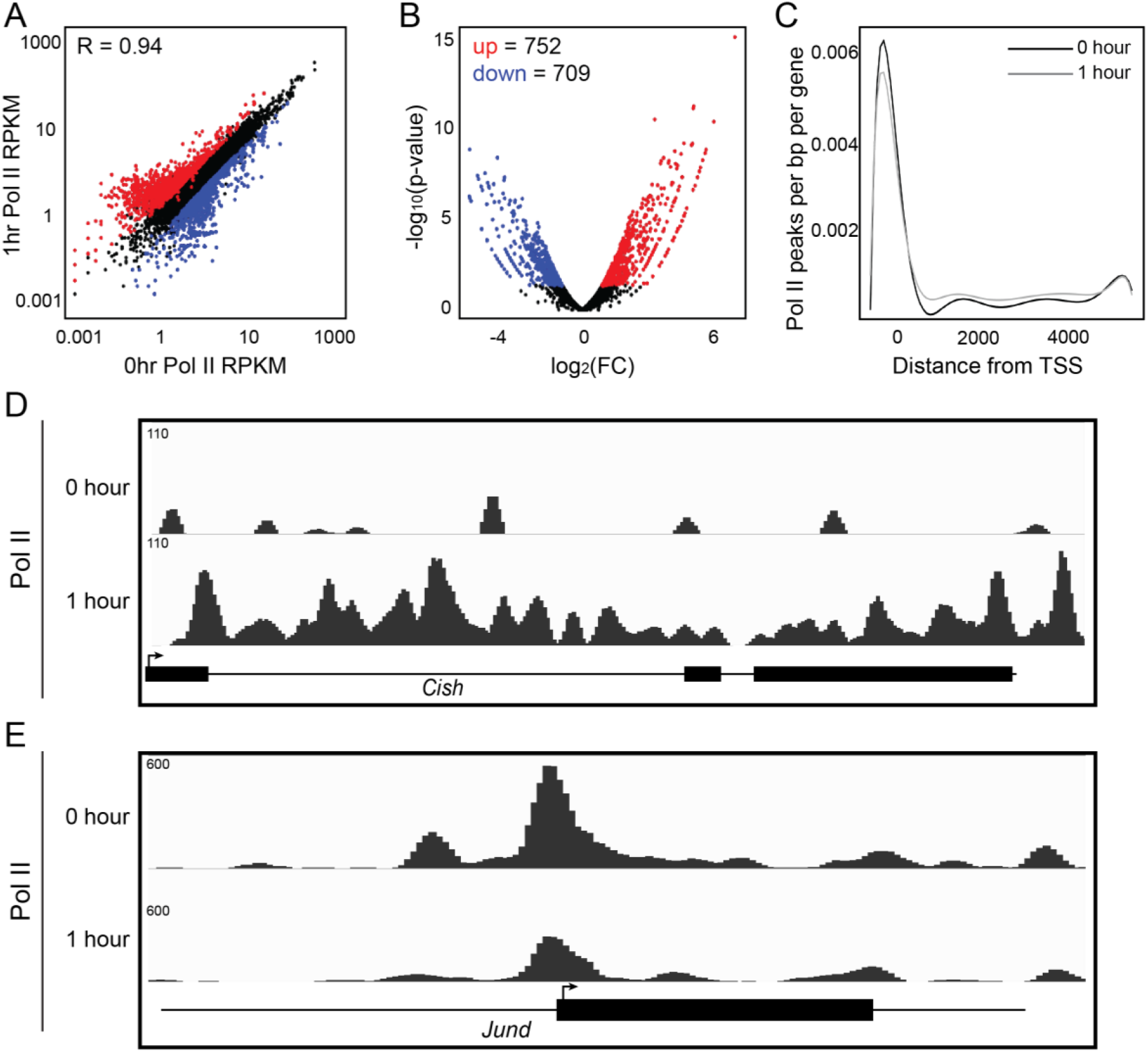
EPO stimulation results in acute transcriptional changes in proerythroblasts. (A) Scatterplot comparing Pol II RPKM before and after 1 hour EPO stimulation. (B) Volcano plot showing significant (p-value < 0.05) differential occupancy of increased (red) and decreased (blue) Pol II after 1 hour EPO stimulation. (C) Metagene plot comparing the position of Pol II peaks relative to transcription start site (TSS) (paired Wilcoxon ranked-sign test, p = 4.882 x 10-11). Genome browser view of ChIP-exo signal for Pol II at the up-regulated *Cish* locus (D) and down-regulated *Jund* locus (E).

**Figure 3.**
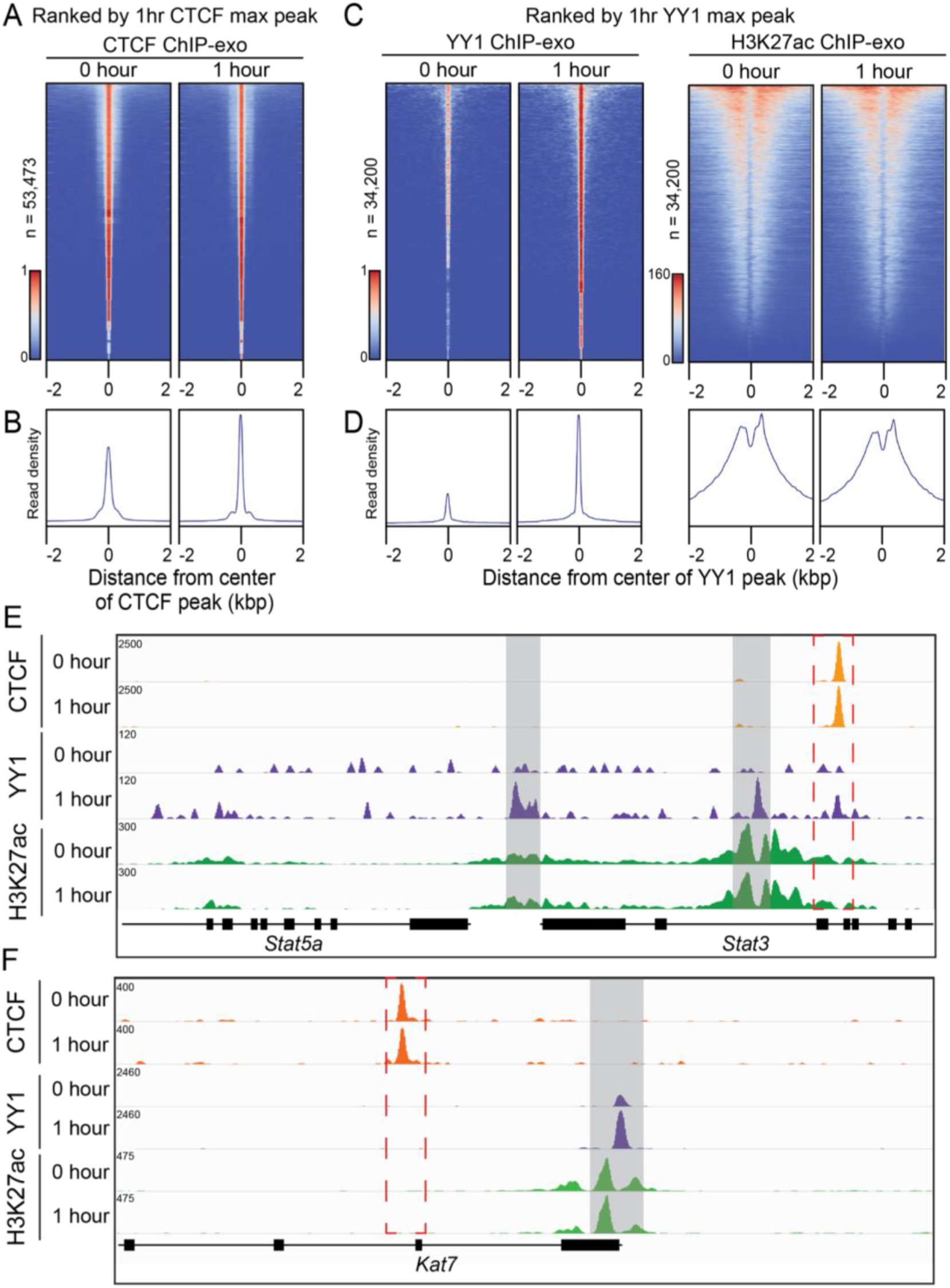
EPO dynamically regulates YY1 occupancy genome-wide. (A) Heatmap of CTCF peaks pre and post EPO stimulation, ranked by 1 hour CTCF max peak. (C) Heatmap of YY1 and H3K27ac peaks pre and post EPO stimulation, ranked by 1 hour YY1 max peak. (B) and (D) Composite plots below each heatmap quantifying the normalized tag density. (E-F) Representative genome browser view of CTCF, YY1, and H3K27ac occupancy in response to EPO stimulation, highlighted in light gray bars and red dashed box.

**Figure 4.**
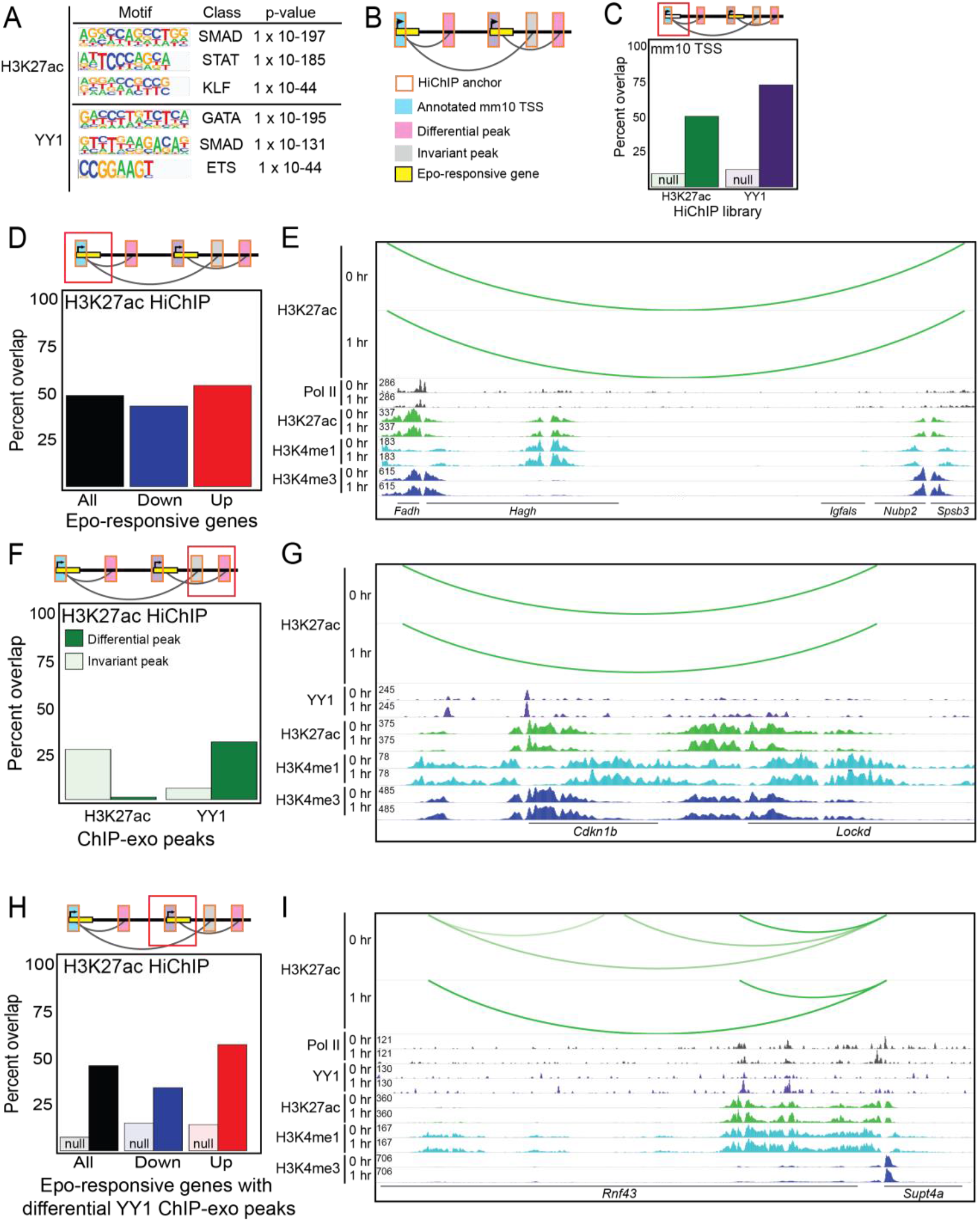
EPO regulates transcription in a pre-established chromatin conformation. (A) TF binding motifs overrepresented in HiChIP loop anchors. (B) A schematic of invariant chromatin landscape. (C) Proportion of interactions with annotated mm10 TSS within their anchor regions. Dark bars represent annotated mm10 TSS and light bars represent randomly generated sequences. (D) Proportion of interactions with promoters of EPO-responsive genes within H3K27ac HiChIP anchor regions. (E) Representative genome browser view of overlap described in (D). (F) Proportion of interactions with differential H3K27ac or YY1 ChIP-exo peaks within anchor regions of H3K27ac HiChIP. Dark bars represent differential ChIP-exo peaks TSS and light bars represent invariant ChIP-exo peaks. (G) Representative genome browser view of overlap described in (F). (H) Proportion of interactions with differential YY1 ChIP-exo peaks at promoters of EPO-responsive genes within anchor regions of H3K27ac HiChIP. Dark bars represent EPO-responsive genes and light bars represent non-responsive genes. (I) Representative genome browser view of overlap described in (H).

## METHODS

### Isolation of Proerythroblasts from FVA infected mice

Highly purified proerythroblasts were obtained from spleens of mice infected with the Friend virus as previously described(Koury et al., 1984; Sawyer et al., 1987), with the following modifications. All animal procedures were performed in compliance with and approval from the Vanderbilt Division of Animal Care (DAC) and Institutional Animal Care and Use Committee (IACUC). Female BALB/cJ mice (12 weeks old, Jackson Laboratories) were infected via intraperitoneal injection of ~104 spleen focus-forming units of Anemia-inducing strain of the Friend virus (FVA). At 13 to 15 days post-infection, the mice were sacrificed and spleens removed. The spleens were homogenized to a single cell suspension by passing the minced spleens through a sterile 100 micron nylon mesh filter into sterile solution of 0.2% bovine serum albumin (BSA) in 1x PBS. The filtrate was then repeatedly pipetted to ensure a single cell suspension. The homogenized spleen cells were size-separated by gravity sedimentation for 4 hours at 4°C in a continuous gradient of 1% to 2% deionized BSA. The sedimentation apparatus consisted of a 25cm diameter sedimentation chamber containing a 2.4L BSA gradient, two BSA gradient chambers containing 1.2L 1% and 2% deionized BSA in 1x PBS, and a cell loading chamber (ProScience Inc.) containing the 50ml cell suspension. After 4 hour sedimentation, cells were collected in 50ml fractions, with proerythroblasts typically enriched in fractions 5-20 of 24 total fractions. Typically about 109 proerythroblasts were obtained from the separation of 1010 nucleated spleen cells (6-7g spleen weight) across three 25cm sedimentation chambers.

### Cell Culture Conditions

To study the effects of erythropoietin (Epo) on terminal erythroid differentiation, FVA-derived proerythroblasts were cultured at 106 cells/ml in Iscove-modified Dulbecco medium (IMDM, Life Technologies #12440043), 30% heat-inactivated fetal bovine serum (Gibco, 26140-079), 1% Penicillin-Streptomycin (Gibco #15140-122), 10% deionized BSA, and 100uM alpha-thioglycerol (MP Biomedicals #155723). Terminal erythroid differentiation of purified proerythroblasts was induced by the addition of 0.4 U/ml human recombinant Epo (10kU/ml Epogen by Amgen, NDC 55513-144-10) to media. At the desired times after the addition of Epo, cells were crosslinked by the addition of 1% formaldehyde for 10 minutes for ChIP analysis and 2% formaldehyde for 20 minutes for HiChIP analysis. Crosslinking was then quenched by the addition of 125mM glycine. Crosslinked cells were collected by centrifugation for 5 minutes at 1,000g at 4°C, washed once with 1x PBS, flash frozen in liquid nitrogen, and stored at −80°C until used. For RNA-seq, cells were removed from culture before crosslinking. Samples were spun for 5 minutes at 1,000g at 4°C and the supernatant was aspirated. Pellets were flash frozen in liquid nitrogen and stored at −80°C until used.

### HiChIP

HiChIP was performed as described(Mumbach et al., 2016) with a few modifications.

#### In Situ Contact Generation

50 million cell pellets were resuspended in 2.5ml ice cold Hi-C Lysis Buffer (10mM Tris HCl, 10mM NaCl, 0.2% NP-40, 1X protease inhibitors (Roche, 04693124001)) and split into 10 million cell amounts. Samples were incubated at 4°C for 30 minutes with rotation. Nuclei were pelleted by centrifugation at 2,500g for 5 minutes at 4°C and washed once with 500ul of ice cold Hi-C Lysis Buffer. After removing supernatant, nuclei were resuspended in 100ul of 0.5% SDS and incubated at 62°C for 10 minutes. SDS was quenched by adding 285ul water and 50ul 10% Triton X-100. Samples were vortexed and incubated for 15 minutes at 37°C. After the addition of 50ul of 10X NEBBuffer 2 (NEB, B7002) and 1ul of MboI restriction enzyme (NEB, R0147), chromatin was digested at 37°C for 1 hour at 700rpm on Thermomixer. Following digestion, MboI enzyme was heat inactivated by incubating the nuclei at 62°C for 20 minutes. To fill in the restriction fragment overhangs and mark the DNA ends with biotin, 52ul of fill-in master mix, containing 15ul of 1mM biotin-dATP (Jena BioScience, NU-835-BIO14-L), 1.5ul of 10mM dCTP (NEB, N044_S), 1.5ul of 10mM dGTP (NEB, N044_S), 1.5ul of 10mM dTTP (NEB, N044_S), and 10ul of 5 U/ul DNA Polymerase I, Large (Klenow) Fragment (NEB, M0210), was added and the tubes were incubated at 37°C for 1 hour at 700rpm on Thermomixer. Proximity ligation was performed by addition of 948ul of ligation master mix, containing 150ul of 10X NEB T4 DNA ligase buffer (NEB, B0202), 125ul of 10% Triton X-100, 15ul of 10 mg/mL BSA (NEB, B9000), 10ul of 400 U/mL T4 DNA ligase (NEB, M0202), and 648ul of water, and incubation at room temperature for 4 hours with rotation.

#### Sonication and Chromatin Immunoprecipitation

After proximity ligation, nuclei were pelleted by centrifugation at 2500g for 5 minutes and resuspended in 880ul Nuclear Lysis Buffer (50mM Tris HCl, 10mM EDTA, 1% SDS, 1X protease inhibitors (Roche, 04693124001)). Samples were vortexed and nuclei were sonicated with a Bioruptor (Diagenode) for 10 minutes on the low setting to solubilize chromatin. Sonicated chromatin was clarified by centrifugation at 16,100g for 15 min at 4°C and supernatant from 10 million cell samples are pooled to a total of 50 million cells. Sample was diluted with 2X ChIP Dilution Buffer (0.01% SDS, 1.1% Triton X-100, 1.2mM EDTA, 16.7mM Tris HCl, 167mM NaCl). 300ul Protein A beads (Thermo, 21348) were washed in 2ml ChIP Dilution Buffer and resuspended in 250ul ChIP Dilution Buffer. Beads were added to 50 million cell sample and incubated at 4°C for 1 hour with rotation. Beads were then separated on a magnetic rack and supernatant was transferred to a new tube. 10ug of antibody for Pol II (Santa Cruz, sc-17798), H3K27ac (Abcam, ab4729), or YY1 (Abcam, ab109237) were added to the tube. Samples were incubated overnight at 4°C with rotation. The next day, 300ul Protein A beads were washed in 2ml ChIP Dilution Buffer and resuspended in 500ul ChIP Dilution Buffer. Beads were added to 50 million cell sample with antibody and incubated at 4°C for 2 hours with rotation. Beads were then separated on a magnetic rack and washed three times with 750ul Low Salt Wash Buffer (0.1% SDS, 1% Triton X-100, 2mM EDTA, 20mM Tris HCl, 150mM NaCl), three times with 750ul High Salt Wash Buffer (0.1% SDS, 1% Triton X-100, 2mM EDTA, 20mM Tris HCl, 500mM NaCl), and three times with 750ul LiCl Wash Buffer (10mM Tris HCl, 250mM LiCl, 1% NP-40, 1% Na-Doc, 1mM EDTA).

#### DNA Elution and Reverse Crosslinking

Beads were then resuspended in 200ul of DNA Elution Buffer (50mM NaHCO_3_, 1% SDS), which is made fresh, and incubated at room temperature for 10 minutes with rotation, followed by 37°C for 3 minutes at 700rpm. Samples were placed on a magnetic rack and supernatant transferred to a new tube. This was repeated once more. 10ul of Proteinase K (Roche, 03115828001) was added to each tube and samples were incubated at 55°C for 45 minutes at 700rpm, followed by 67°C for 1.5 hours at 700rpm. DNA was then purified using Zymo DNA Clean and Concentrator (Zymo, D4003) according to manufacturer’s protocol and eluted in 10ul water. The amount of eluted DNA was quantified by Qubit dsDNA HS kit (Invitrogen, Q32854).

#### Biotin Capture and Sequencing Preparation

25ul of Streptavidin C-1 beads (Invitrogen, 65001) were washed with 1ml Tween Wash Buffer (5MM Tris HCl, 0.5mM EDTA, 1M NaCl, 0.05% Tween-20) and resuspended in 10ul of 2X Biotin Binding Buffer (10mM Tris HCl, 1mM EDTA, 2M NaCl). 10ul of bead mixture was added to 50ng of purified DNA for each sample, incubating at room temperature for 15 minutes, agitating every 5 minutes. After capture, beads were separated with a magnet and the supernatant was discarded. Beads were then washed twice with 500ul of Tween Wash Buffer, incubating at 55°C for 2 minutes at 700rpm. Beads were washed with 100ul 1X TD Buffer (diluted from 2X TD Buffer (20mM Tris HCl, 10mM MgCl_2_, 20% Dimethylformamide)). Beads were resuspended in 50ul of master mix, containing 25ul 2X TD Buffer, 2.5ul Tn5 Tagment DNA enzyme (Illumina, 15027865), and 22.5ul water. Samples were incubated at 55°C for 10 minutes at 700rpm. Beads were separated on a magnet and supernatant was discarded. Beads were washed with 750ul of 50mMEDTA at 50°C for 30 minutes, washed twice with 750ul of 50mMEDTA at 50°C for 3 minutes each, then washed twice with 750ul of Tween Wash Buffer at 55°C for 2 minutes each, and finally washed once with 750uL of 10mM Tris HCl pH 7.5. Beads were separated on a magnet and supernatant was discarded.

#### PCR and Size Selection

To generate the sequencing library, PCR amplification of the tagmented DNA was performed while the DNA is still bound to the beads. Beads were resuspended in a PCR master mix, consisting of 36ul water, 1.25 unique Nextera Ad2.X primer, 10ul Phusion HF 5X buffer (NEB, E0553), 1ul 10mM dNTPs, 1.25ul universal Nextera Ad1 primer, and 0.5ul Phusion DNA Polymerase (NEB, E0553). DNA was amplified with 8 cycles of PCR. After PCR, beads were separated on a magnet and the supernatant containing the PCR amplified library was transferred to a new tube, purified using the Zymo DNA Clean and Concentrator (Zymo D4003) kit according to manufacturer’s protocol and eluted in 52ul water. Purified HiChIP libraries were size selected to 300-700 basepairs using a double size selection with AMPure XP beads (Beckman Coulter, A68831). HiChIP libraries were paired-end sequenced on an Illumina NextSeq500 with reads 75 nucleotides in length.

### Chromatin Immunoprecipitation with Lambda Exonuclease Digestion (ChIP-exo)

With the following modifications, ChIP-exo was performed as previously described(Perreault and Venters, 2016; Rhee and Pugh, 2011) with chromatin extracted from 50 million cells, ProteinG MagSepharose resin (GE Healthcare), and 10ug of antibody directed against Pol II (Santa Cruz, sc-17798), YY1 (Abcam, ab109237), or CTCF (Millipore, 07-729). First, formaldehyde crosslinked cells were lysed with buffer 1 (50mM HEPES–KOH pH 7.5, 140mM NaCl, 1 mM EDTA, 10% Glycerol, 0.5% NP-40, 0.25% Triton X-100), washed once with buffer 2 (10mM Tris HCL pH 8, 200mM NaCl, 1mM EDTA, 0.5mM EGTA), and the nuclei lysed with buffer 3 (10mM Tris HCl pH 8, 100mM NaCl, 1mM EDTA, 0.5mM EGTA, 0.1% Na–Deoxycholate, 0.5% *N*-lauroylsarcosine). All cell lysis buffers were supplemented with fresh EDTA-free complete protease inhibitor cocktail (CPI, Roche #11836153001). Purified chromatin was sonicated with a Bioruptor (Diagenode) to obtain fragments with a size range between 100 and 500 base pairs. Triton X-100 was added to extract at 1% to neutralize sarcosine.

Insoluble chromatin debris was removed by centrifugation, and sonication extracts stored at −80°C until used for ChIP analysis. Libraries were sequenced using an Illumina NextSeq500 sequencer as single-end reads 75 nucleotides in length on high output mode. To assess reproducibility of biological replicates, Pearson’s correlation was calculated (Supp. Fig. 1).

### RNA-seq

RNA was isolated using the Qiagen RNAeasy kit (Qiagen, 74104) per manufacturer’s instructions. Stranded polyA selected libraries were prepared using NEBNext PolyA mRNA isolation standard protocol, NEBNext rRNA Depletion standard protocol, and finally NEBNext Ultra II Directional DNA library preparation kit (Illumina, E75530S) per manufacturer’s protocol. PCR amplified RNA-seq libraries were size selected using AMPure XP beads (Beckman Coulter, A68831). RNA-seq libraries were subjected to 75 basepair single end sequencing on Illumina NextSeq500 sequencer.

### HiChiP data analysis

#### Alignment

HiChIP library sequence reads were aligned to the mouse mm10 reference genome using HiC-Pro(Servant et al., 2015) with the following options in the configuration file:

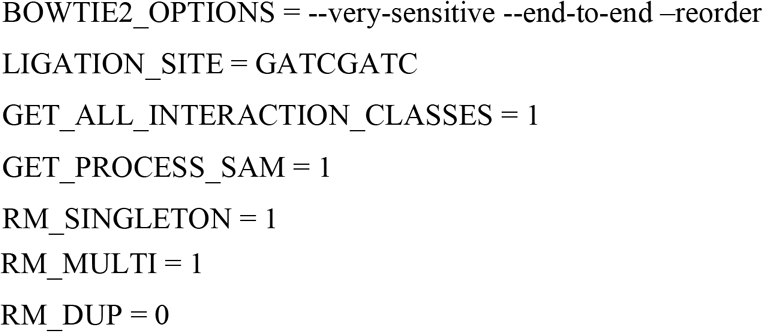

#### Chromatin interaction identification

Hichipper(Lareau and Aryee, 2018b) was applied to HiC-Pro output files to identify high confidence chromatin contacts using EACH, ALL peak finding settings. The quickAssoc and annotateLoops functions in the diffloop R package(Lareau and Aryee, 2018a) were used to find differential loops and annotate epigenetic features, respectively. Enhancers were denoted as the intersection of H3K4me1 and H3K27ac peaks (previously published data, Key Resources Table) and promoters were identified using the getMouseTSS function.

#### HiChIP display

To visualize chromatin interactions identified using HiChIP, the -make-ucsc option was added when analyzing the data using hichipper(Lareau and Aryee, 2018b).

### ChIP-exo data analysis

#### Alignment

ChIP-exo library sequence reads were aligned to the mouse mm10 reference genome using BWA-MEM algorithm(Li and Durbin, 2010) using default parameters. The resulting bam files were first sorted using the Samtools Sort function(Li et al., 2009), and then bam index files were generated using the Samtools Index function(Li et al., 2009). Since patterns described here were evident among individual biological replicates, and replicates were well correlated, we merged all tags from biological replicate data sets for final analyses.

#### Peak calling

ChIP-exo peaks were annotated and quantified using the Hypergeometric Optimization of Motif EnRichment (HOMER) suite(Heinz et al., 2010). Briefly, bam files were converted to tag directories using the makeTagDirectory function with the –genome, –checkGC, and –format options. The findPeaks function was used to identify ChIP peaks using –o auto and –style gro-seq or factor for Pol II or CTCF/YY1 libraries, respectively. To quantify and normalize tags to RPKM, the analyzeRepeats function was used with the –rpkm, –count genes, –strand both, –condenseGenes, and –d options.

#### Heatmaps

bigWig files for CTCF and YY1 libraries were generated using the deepTools bamCoverage function(Ramirez et al., 2014). To create aligned heatmaps, first a matrix was generated using the computeMatrix function with the following options: reference-point –S, –a 2000, –b 2000, –referencePoint center, –verbose, –missingDataAsZero, and –p max/2. Then, the heatmap was created using the plotHeatmap function with the following options: –verbose and –sortRegions descend.

#### ChIP-exo display

Raw sequencing tags were smoothed (20 basepair bin, 100 basepair sliding window) and normalized to reads per kilobase per million (RPKM) using deepTOOLS(Ramirez et al., 2014) and visualized with Integrative Genomics Viewer (IGV)(Robinson et al., 2011).

### RNA-seq data analysis

#### RNA-seq alignment, transcript assembly, and differential expression

RNA-seq library sequence reads were aligned to the mouse mm10 reference genome using TopHat(Trapnell et al., 2009) using default parameters. Cufflinks(Trapnell et al., 2012) was used to assemble transcripts and quantify expression of transcripts. Cuffmerge(Trapnell et al., 2012) merges all transcript assemblies to create a single merged transcriptome annotation for final analyses.

#### RNA-seq display

CummeRbund visualizes RNA-seq data analyzed using cufflinks.

### Definition of regulatory regions

Throughout the manuscript multiple analyses rely on overlaps with different regulatory regions, namely enhancers and promoters. Here we explain how these regulatory regions were defined.

**Promoters** are defined here as the comprehensive list of annotated transcription start sites (TSS) in the mm10 mouse genome from UCSC.

**Enhancers** are defined here as regions of the genome marked by H3K4me1 and H3K27ac. This group is further supported by enhancer identified using ChromHMM in (Perreault et al., 2017).

### Definition of chromatin features

Throughout the manuscript multiple analyses rely on overlaps with different chromatin features. Here we explain how these features were defined.

**HiChIP anchors**, as identified by the hichipper analysis pipeline, are the regions between restriction enzyme motifs that contain a ChIP peak for the factor of interest. In the present study, we use anchor as a broad term to define the endpoints of a HiChIP loop.

**HiChIP loops** are chromatin interactions that occur between two anchors. These loops have a specific score, which is the number of paired-end tags (PETs) that support the interaction.

**Invariant loops** are chromatin interactions that have a fold change in score (as calculated by diffloop) to be between −2 and 2.

### Data and Code Availability

Raw and preprocessed data will be available once this manuscript it accepted for publication.

## RESULTS

### EPO induces terminal erythrocyte differentiation through changes in gene expression

The anemia-inducing strain of the Friend Virus (FVA) system enables us to investigate the temporal dynamics of gene regulation and genome architecture in response to hormone stimulation. In this model, systemic treatment of mice with FVA induces ProEB proliferation in the spleen. After 14 days, large quantities of lineage committed ProEBs can be isolated and purified. Stimulation of ProEBs with the hormone EPO in an *ex vivo* culture system induces synchronous terminal differentiation into mature erythrocytes over a 48 hour period (Fig. 1A) (Sawyer et al., 1987).

Using this model system, we observed a predictable shift in size and shape of maturing erythroid precursors during erythropoiesis, as visualized by light microscopy of Haemotoxylin and Eosin stained cells. Before purification, the cells appear heterogeneous (Fig. 1B, Input). After purification, a uniform population of ProEBs is obtained, evident as large, round cells. This morphological stage persists until approximately 12 hours after the start of EPO stimulation (Fig. 1B). After 24 hours of EPO, cells form polychromatic erythroblasts (PolyEBs), characterized by the accumulation of hemoglobin, which coincides with the continued increase in globin gene expression. Finally, after 48 hours of *ex vivo* culture in EPO, the cells have terminally differentiated into reticulocytes, marked by high hemoglobin production and nucleus extrusion (Fig. 1B, 48hr).

Having established the kinetics of this model system, we next examined how differentiation impacts the erythroid gene expression program over time. RNA-sequencing (RNA-seq) at 6 time points demonstrated both up-regulation and down-regulation of genes as compared to EPO-naïve cells (Fig. 1C). Overall, approximately 12,000 genes had differential expression during the entire 48 hour time course of erythropoiesis, with 8,105 of those genes significantly differentially expressed (p < 0.05). More specifically, there were 439 genes that significantly changed expression after only 1 hour of EPO (p < 0.05). These transcriptomic data highlight the large changes in gene expression that accompany the morphological shifts occurring during erythropoiesis.

### EPO activates rapid transcriptional changes in ProEBs

The progressive changes in gene expression observed by RNA-seq reflect global transcriptional responses. We set out to investigate the immediate transcriptional response to EPO in purified ProEBs. To assess the acute effect of hormone stimulation on transcription, we performed ChIP-exo for RNA polymerase II (Pol II) before and after 1 hour of EPO exposure. Overall, Pol II occupancy is highly correlated when comparing ChIP-exo signal pre- and post-EPO stimulation in this short time frame (Fig. 2A). This result indicates that global Pol II occupancy does not change at the majority of transcribed genes after 1 hour. However, analysis of fold change of Pol II signal in these conditions did identify significant differential occupancy of Pol II at a smaller subset of genes (p < 0.05) (Fig. 2B). We detected both significantly increased (n = 752, red) and decreased (n = 709, blue) Pol II signal at these EPO-responsive loci, indicating a set of genes that are regulated at the transcriptional level by EPO after only 1 hour.

Rapid transcriptional induction may reflect PIC assembly, pause release of Pol II, or a combination of these steps. To investigate the mechanism of EPO-regulated transcription, we next mapped Pol II signal at transcriptional start sites (TSS) or gene bodies of all induced genes. This approach identified that Pol II is more abundant at the TSS of genes before EPO (Fig. 2C). 1 hour after EPO stimulation, Pol II transitions beyond the TSS into the gene body, indicative of pause release of Pol II at induced genes. The dynamics of increased Pol II occupancy can be visualized at an exemplary locus of cytokine inducible SH2 containing protein (*Cish*) (Fig. 2D). *Cish* is a known target of the JAK-STAT signaling pathway, which is directly activated by EPO(Matsumoto et al., 1997; Rascle and Lees, 2003).

We also observed down-regulation of transcription, including the *Jund* gene (Fig. 2E). *Jund* is a component of the AP1 complex, which regulates response to cytokines, growth factors, stress, and infections in a variety of cellular contexts(Hernandez et al., 2008; Karin et al., 1997). *Jund*, along with other members of the Jun family, has been found to prevent differentiation in murine erythroleukemic cells(Prochownik et al., 1990), highlighting the critical need to down-regulate this gene during erythropoiesis. The importance of *Cish* and *Jund* in differentiation provide specific examples of the biological significance of the early EPO-mediated transcriptional responses described.

### EPO dynamically regulates YY1 occupancy genome-wide

Signal dependent activation of Pol II is accompanied by alterations in chromatin organization. However, the impact of EPO on chromatin structure and function during erythropoiesis is not well understood. To begin addressing this question, we first examined the genome-wide occupancy patterns of CTCF and YY1, two TFs known to play key roles in genome organization and gene regulation. CTCF occupancy did not change between pre and post 1 hour EPO treatment, as demonstrated by the comparison of global enrichment analysis at each time point (Fig. 3A, B). The absence of changes in CTCF binding reflect the fact that the cells in each treatment group are lineage committed ProEBs (Fig. 1A, B). Thus, the invariant occupancy of CTCF observed is consistent with recent studies demonstrating CTCF decreases variability in gene expression and thereby functions to maintain an established cell state(Ren et al., 2017).

In contrast to CTCF, YY1 is rapidly redistributed in the genome following 1 hour EPO stimulation (Fig. 3C, D). In EPO-naïve ProEBs, the majority of YY1 localized to intergenic regions (48%, Supp. Fig. 2A). However, this distribution shifted to intronic sites after EPO (42%, Supp. Fig. 2A). Notably, YY1 ChIP-exo signal at TSSs significantly increased from 5% to 17% following EPO (Supp. Fig. 2A). Comparison of CTCF and YY1 also revealed minimal overlap in localization of these two factors (Supp. Fig. 2B, 7% and 5%, respectively); and EPO did not alter the proportion of shared regions (Supp. Fig. 2B). In addition, ranking of H3K27ac signal by YY1 enrichment demonstrated that H3K27ac signal did not change in an appreciable manner compared to YY1 after EPO. This suggests that the subset of YY1 sites were more dynamic than H3K27ac, which is commonly used to identify active enhancers in the genome (Fig. 3C, D). These results suggest that the chromatin domains established by CTCF and YY1 are distinct and these structural proteins have unique functions during EPO-dependent gene regulation in ProEBs.

At the *Stat3* gene, an exemplary locus of EPO-mediated transcription, a strong CTCF peak is evident pre-EPO and does not change after EPO (Fig. 3E). At the same *Stat3* locus, YY1 occupancy increases at multiple *de novo* binding sites, as well as one site that overlaps with CTCF. Similar changes in YY1 can be visualized at another EPO-responsive gene, *Kat7* (Fig. 3F). These specific loci illustrate how EPO induces a dynamic change in YY1 occupancy at a subset of genes relevant to signal transduction and chromatin modification during erythropoiesis.

### EPO regulates transcription in a pre-established chromatin conformation

Our discovery that YY1 rapidly redistributes in the ProEB genome following EPO stimulation prompted us to explore the role of YY1 and H3K27ac in chromatin organization. To accomplish this goal, we examined the global chromatin contacts mediated by YY1 or H3K27ac in EPO-stimulated ProEBs using HiChIP. HiChIP is a chromosome conformation capture assay that maps chromatin interactions between specific factors genome-wide (Mumbach et al., 2016). Using the hichipper program(Lareau and Aryee, 2018b), loops are defined as the distance between two ends of a chromatin interaction called anchors, which are identified in hichipper by extending ChIP peaks to the edges of the restriction fragment (Supp. Fig. 5). As a consequence of this computational definition, HiChIP anchors typically span a wide range of lengths. In our dataset, H3K27ac and YY1 anchors were approximately 4 kilobases (kb) on average both before and after EPO (Supp. Fig. 6A). In addition, the average loop lengths for interactions in either H3K27ac or YY1 anchors were approximately 317 kb, with no evident change in loop length in response to EPO treatment (Supp. Fig. 6B).

To investigate the biological significance of these loops in ProEBs, we first conducted unbiased, *de novo* motif discovery analysis using DNA sequences from all HiChIP anchor regions. Strikingly, we identified an enrichment of consensus motifs of multiple TFs known to regulate specification of the erythroid lineage, including STAT(Kisseleva et al., 2002; Watowich, 2011), KLF(Cantor and Orkin, 2002; Kang et al., 2015; Miller and Bieker, 1993), and GATA(Cantor and Orkin, 2002; Lentjes et al., 2016; Weiss and Orkin, 1995) (Fig. 4A). Additionally, auxiliary factors, such as SMAD and ETS, aid in the maintenance of gene expression and lineage commitment, respectively(Pimkin et al., 2014; Schmerer and Evans, 2003; Schuetze et al., 1993). The enrichment of these consensus motifs within YY1 and H3K27ac HiChIP anchor regions suggests functional coupling between erythroid TFs and chromatin conformation during erythropoiesis.

After defining the general characteristics of H3K27ac- and YY1-mediated interactions, we next determined whether genome architecture changes in response to EPO stimulation. Overall, we identified 151,032 H3K27ac- and 138,210 YY1-mediated chromatin contacts in cells pre- and post-EPO treatment using diffloop(Lareau and Aryee, 2018a). In hichipper and diffloop, these interactions are ranked by loop score, which is defined as the number of paired end tags that support a given interaction. The majority of these loops had a score less than 5 (Supp. Fig. 6C, 6D), indicating that weak interactions predominate in both conditions. Moreover, using loop scores as a metric for chromatin organization, we were not able to detect any significant change in the overall chromatin interactions in response to EPO. These data suggest that acute EPO-induced transcriptional regulation occurs within an invariant chromatin architecture in lineage-committed ProEBs.

Based on these data, we then focused on invariant loops, defined by fold change in loop score between −2 and 2, to gain insight into the relationship between YY1- and H3K27ac-mediated HiChIP loops and EPO-dependent transcription. We reasoned that a better understanding of the interactions between E-P sites might provide new insights into how EPO regulates transcription. First, we quantified the proportion of HiChIP anchors (H3K27ac or YY1) of invariant E-P loops that map to UCSC annotated TSSs in the mm10 reference genome (Fig. 4B). We would expect 50% of the anchors to be found in TSS regions if each promoter was connected to one enhancer. Indeed, half of the anchors of chromatin interactions mediated by H3K27ac were observed in annotated TSSs (50%, Fig. 4C). In contrast to H3K27ac, 73% of chromatin interactions with YY1 anchors overlapped with annotated TSSs, indicating that more YY1 anchors are at promoters compared to enhancer regions (Fig. 4C). For comparison, we conducted a test of the null hypothesis by overlapping HiChIP anchors with randomly generated DNA regions. This analysis resulted a low percent overlap for both H3K27ac (9%) and YY1 (12%) under these conditions (Fig. 4C, light bars). These results suggest that a single enhancer could regulate the transcription of multiple target genes in chromatin interactions mediated by YY1.

Given that E-P interactions identified by HiChIP overlap with annotated promoter regions, we next examined the relationship between these loops and the associated changes in transcriptional response to EPO, as described in Figure 2. Focusing only on EPO-induced genes (n = 1,462), we identified a higher proportion of overlap for H3K27ac anchors at promoters of EPO-responsive genes as compared to YY1 anchors (50% vs 6% respectively, Fig. 4D, Supp. Fig. 7A). We also examined this relationship as a function of genes induced (n=752) or down-regulated (n=709) by EPO. As expected, fewer promoters in EPO down-regulated genes overlapped with H3K27ac anchors. The persistence of H3K27ac in down-regulated promoters is consistent with prior work demonstrating that loss of H3K27ac signal enhancers and promoters can lag behind a decrease in transcription(Brown et al., 2014). Figure 4E shows a representative example of this overlap at the *Fadh1* gene, which is intricately involved in mitochondrial activity and metabolism(Weiss et al., 2018). Mitochondrial biogenesis is activated by EPO and is therefore highly regulated during erythropoiesis(Carraway et al., 2010; Liu et al., 2017). These data reveal that H3K27ac-mediated loops are strongly connected to EPO-induced transcriptional response.

We were surprised that H3K27ac anchors overlapped at promoters of EPO-induced genes more than YY1 anchors, given that YY1 genome occupancy measured by ChIP-exo was more dynamic after EPO treatment (Fig. 3). To resolve this apparent paradox, we first examined the relationship between differential H3K27ac or YY1 occupancy measured with ChIP-exo and all invariant chromatin loops identified by HiChIP. With this approach, we detected significant enrichment for differential YY1 ChIP-exo peaks at invariant H3K27ac anchors (Fig. 4F). In contrast to this result, invariant H3K27ac loops were associated with invariant H3K27ac ChIP-exo peaks (Fig. 4F). An example of this relationship can be observed at the *Cdkn1b and Lockd* loci, two genes that regulate exit of erythroid precursors from the cell cycle, a required step in differentiation(Paralkar et al., 2016) (Fig. 4G). Anchors of invariant YY1 chromatin loops were also enriched at loci with differential YY1 ChIP-exo peaks (Supp. Fig. 7B).

Analysis of the data focusing only on EPO-responsive genes revealed that H3K27ac anchors remained more enriched at promoters of EPO-responsive genes with differential YY1 ChIP-exo peaks (Fig. 4H). By contrast, YY1 anchors were not enriched at promoters to the same degree as regions of differential YY1 chromatin occupancy measured by ChIP-exo (Fig. 4H, Supp. Fig. 7C). A representative example of this overlap is shown at the *Supt4a* gene, which encodes the SPT4 protein, a component of the DSIF elongation complex(Crickard et al., 2017; Schneider et al., 2006), implicating this locus in transcriptional regulation. Together, these results support a model whereby EPO induces dynamic transcription and TF binding within a pre-established chromatin context.

## DISCUSSION

The findings presented here examining erythrocyte differentiation in response to EPO are consistent with an emerging paradigm that signal-dependent transcriptional responses occur within a pre-established chromatin landscape identified using HiC methodologies. For example, TNFalpha-responsive enhancers in human fibroblasts were already in contact with their target promoters before signaling. These results suggest a model in which signal-responsive TFs bind to enhancers to function within a chromatin architecture that is pre-established(Jin et al., 2013). Glucocorticoid treatment in human A549 cells revealed that glucocorticoid receptor binding to the genome did not promote new chromatin contacts, but instead induced changes in existing interactions to regulate transcription(D’Ippolito et al., 2018). HiC analysis in Drosophila S2 and human K562 cells identified that no global changes in TADs emerged after heat shock treatment, despite changes in TF binding and induction of heat shock response genes(Ray et al., 2019). Finally, capture HiC and 4C experiments in ESCs have provided evidence that hardwired chromatin interactions provide an environment for TF binding and enhancer activation that facilitates a rapid transcriptional response to signaling in neuronal development(Atlasi et al., 2019). Unlike these studies, our study employed the HiChIP assay to define the genome-wide contacts mediated by specific factors, namely H3K27ac and YY1.These H3K27ac and YY1 HiChIP contacts revealed a subset of invariant chromatin loops that connect enhancers and EPO-regulated genes, thereby refining the E-P connectome in erythroid cells. These chromatin interactions provide novel insights to conformational features, such as enhancer skipping and promoter-promoter interactions, which cannot be determined using 1D chromatin features(Mumbach et al., 2017). Future work will investigate these confirmation features to evaluate previously identified E-P interactions in ProEBs(Perreault et al., 2017).

Given that CTCF domains shift during development(Nora et al., 2017), we originally hypothesized that EPO would induce changes to CTCF occupancy and subTAD organization. However, CTCF occupancy did not change but instead decreased after EPO, supporting the idea of selective pruning of CTCF binding sites during differentiation. This idea was described in neuronal development(Beagan et al., 2017). In contrast, YY1 did redistribute dynamically in the genome within 1 hour of EPO stimulation, suggesting a more critical role for YY1 in chromatin organization in early erythroid maturation. These data are consistent with studies identifying YY1’s role in E-P loops and transcriptional activation(Weintraub et al., 2017). YY1 occupancy was also more dynamic than H3K27ac, suggesting that YY1 chromatin occupancy may improve assignment of enhancers to target genes. We speculate that combined H3K27ac and YY1 occupancy data may assign enhancers to target genes with more accuracy than H3K27ac alone, given that EPO had a much larger effect on YY1 in ProEBs. This concept will require additional studies.

Not surprisingly, we did observe dynamic changes in Pol II occupancy in response to EPO at a subset of genes both significantly up- and down-regulated. These results are consistent with a growing body of work identifying paused Pol II at signal-responsive genes. It has been proposed that this state of Pol II enables rapid transcriptional response to environmental stimuli. For example, in Drosophila S2 cells stalled Pol II was strongly enriched at genes that are induced by multiple signaling pathways involved in regulating development, cell differentiation, and cell communication(Muse et al., 2007). Additionally, study of murine macrophage cell lines identified an accumulation of paused Pol II at the *TNFalpha* gene in quiescent cells before induction of the gene by inflammatory cytokines(Adelman et al., 2009).

There are still several aspects of EPO’s impact on transcription and chromatin structure that remain unanswered. We identified discordance between dynamic YY1 binding measured by ChIP-exo and invariant YY1-mediated interactions determined by HiChIP. The majority of YY1 HiChIP loops had weak scores (paired-end tags < 5, Supp. Fig. 6), despite abundant YY1 binding in the genome. This suggests that the overall abundance of YY1 does not necessarily indicate the strength of the loop it mediates. It is possible that YY1 binding locations are establishing chromatin contacts that will gain strength over time, and therefore delineate cell-type specific interactions more decisively as maturation continues. Additionally, we expected Pol II ChIP-exo differential peaks to be found at gene promoters that exhibited differential expression as measured by RNA-seq. However, we only detected a small overlap in the gene promoters where this was the case. It is likely that steady-state gene expression measured by RNA-seq lags behind rapid transcriptional responses assessed by Pol II ChIP-exo. Future studies will investigate this relationship between Pol II occupancy and gene expression across the entire period of erythrocyte maturation in the FVA model.

Together, the results presented here integrate epigenome and transcriptional profiles with genome-wide HiChIP datasets to describe how hormone stimulation regulates erythroid differentiation. We demonstrate that chromatin domains are not as dynamic as these other features. Future work will focus on integrating changes in Pol II, CTCF, H3K27ac, and YY1, as well as the chromatin contacts they mediate, during erythropoiesis with the goal of understanding how the 3D genome influences transcription and dynamic gene regulatory programs during erythrocyte maturation.

## Supplemental Figure Legends

**Supplemental Figure 1.**
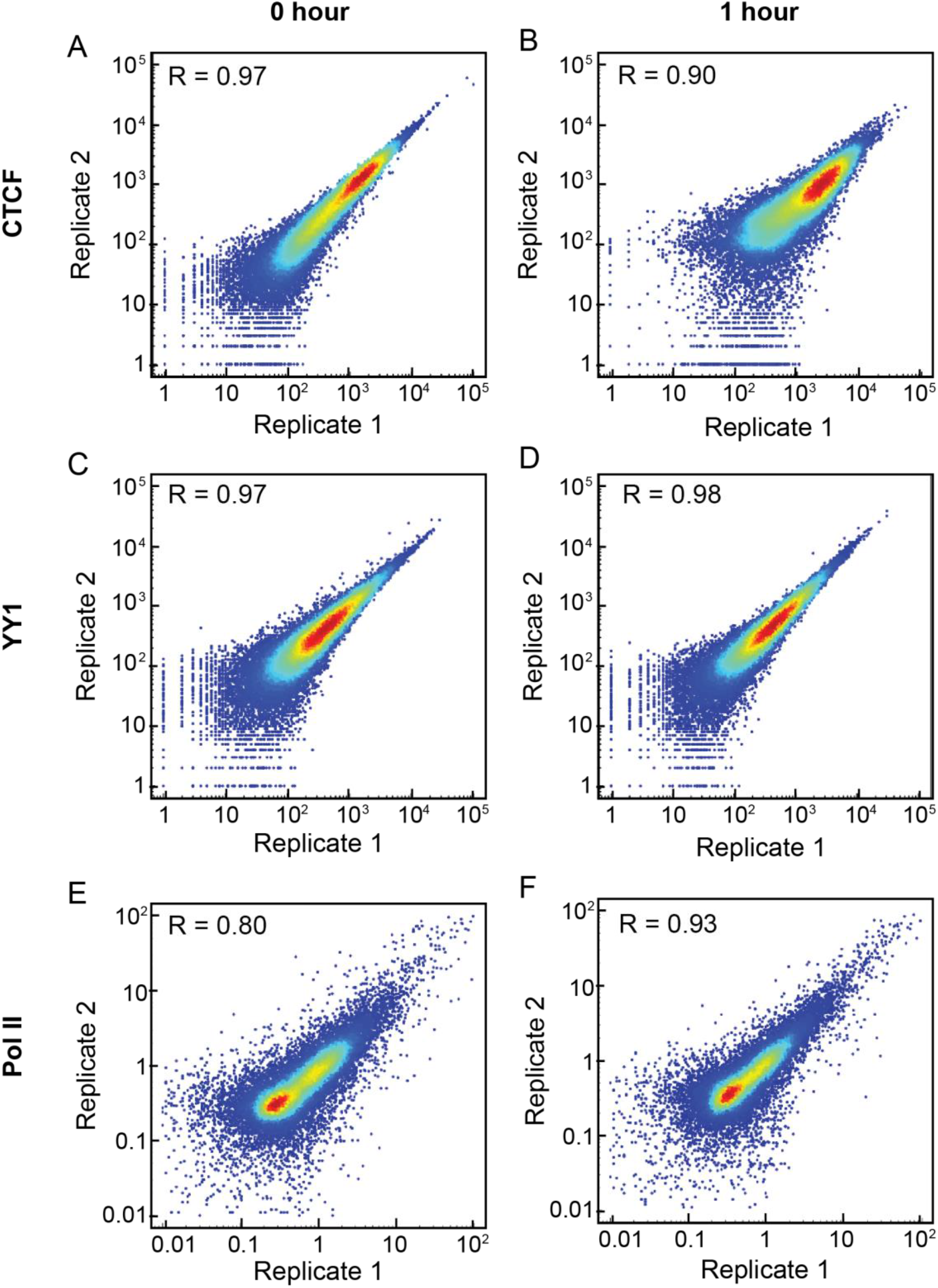
Biological replicate correlation for ChIP-exo libraries. Related to Figures 2 and 3. (A) Biological replicates of peaks called at 0 hour Epo for CTCF. Scatterplot is displayed in log scale, with Pearson’s correlation coefficient in the upper left corner. (B) Biological replicates of peaks called at 1 hour Epo for CTCF. Scatterplot is displayed in log scale, with Pearson’s correlation coefficient in the upper left corner. (C) Biological replicates of peaks called at 0 hour Epo for YY1. Scatterplot is displayed in log scale, with Pearson’s correlation coefficient in the upper left corner. (D) Biological replicates of peaks called at 1 hour Epo for YY1. Scatterplot is displayed in log scale, with Pearson’s correlation coefficient in the upper left corner. (E) Biological replicates of peaks called at 0 hour Epo for Pol II. Scatterplot is displayed in log scale, with Pearson’s correlation coefficient in the upper left corner. (F) Biological replicates of peaks called at 1 hour Epo for Pol II. Scatterplot is displayed in log scale, with Pearson’s correlation coefficient in the upper left corner.

**Supplemental Figure 2.**
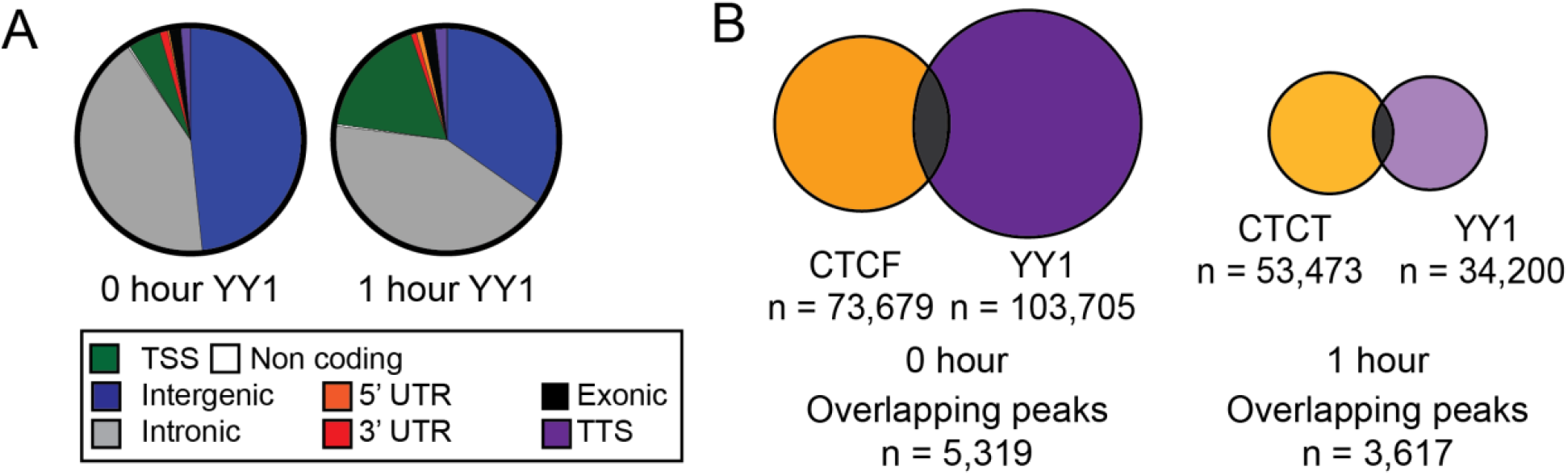
Epo dynamically regulates YY1 occupancy genome-wide. Related to Figure 3. (A) YY1 binding locations in the genome. (B) Comparison of CTCF and YY1 peak overlap before and after 1 hour Epo stimulation.

**Supplemental Figure 3.**
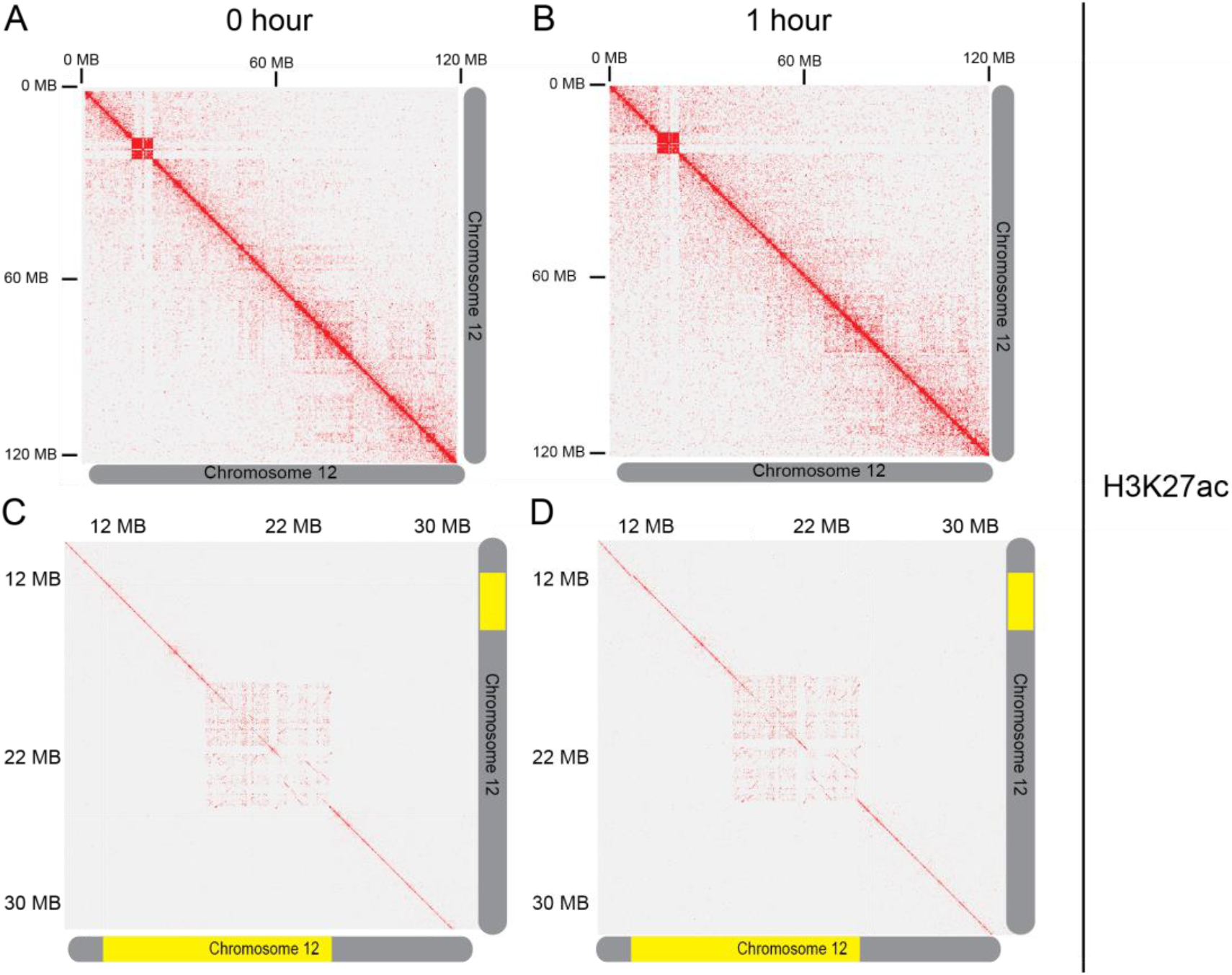
Chromatin contact maps for H3K27ac HiChIP. Related to Figure 4. (A) Chromatin contacts at 0 hour Epo on chromosome 12 at 250KB resolution. (B) Chromatin contacts at 1 hour Epo on chromosome 12 at 250KB resolution. (C) Chromatin contacts at 0 hour Epo on chromosome 12 at 25KB resolution. (D) Chromatin contacts at 1 hour Epo on chromosome 12 at 25KB resolution.

**Supplemental Figure 4.**
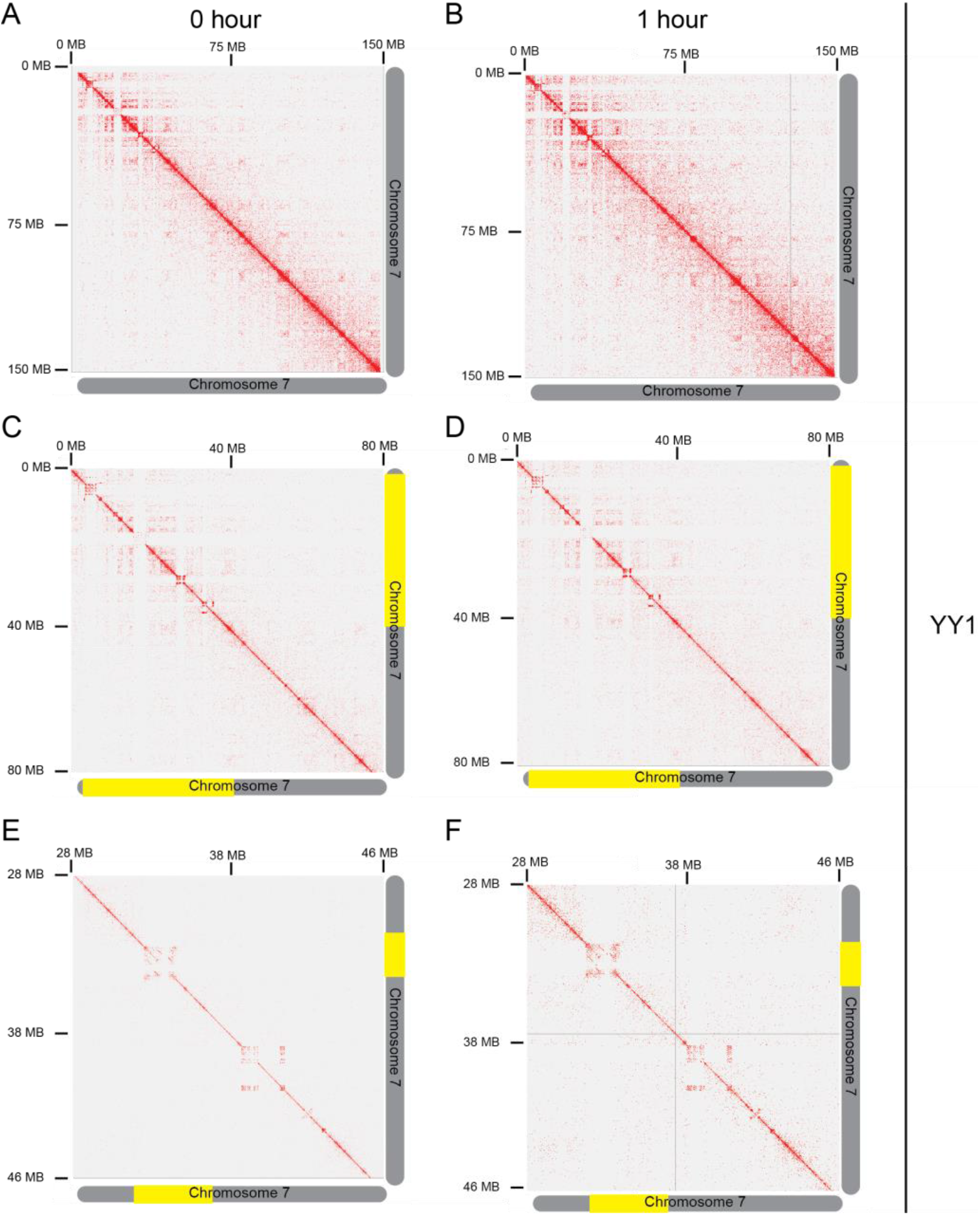
Chromatin contact maps for YY1 HiChIP. Related to Figure 4. (A) Chromatin contacts at 0 hour Epo on chromosome 7 at 250KB resolution. (B) Chromatin contacts at 1 hour Epo on chromosome 7 at 250KB resolution. (C) Chromatin contacts at 0 hour Epo on chromosome 7 at 100KB resolution. (D) Chromatin contacts at 1 hour Epo on chromosome 7 at 100KB resolution. (E) Chromatin contacts at 0 hour Epo on chromosome 7 at 25KB resolution. (F) Chromatin contacts at 1 hour Epo on chromosome 7 at 25KB resolution.

**Supplemental Figure 5.**
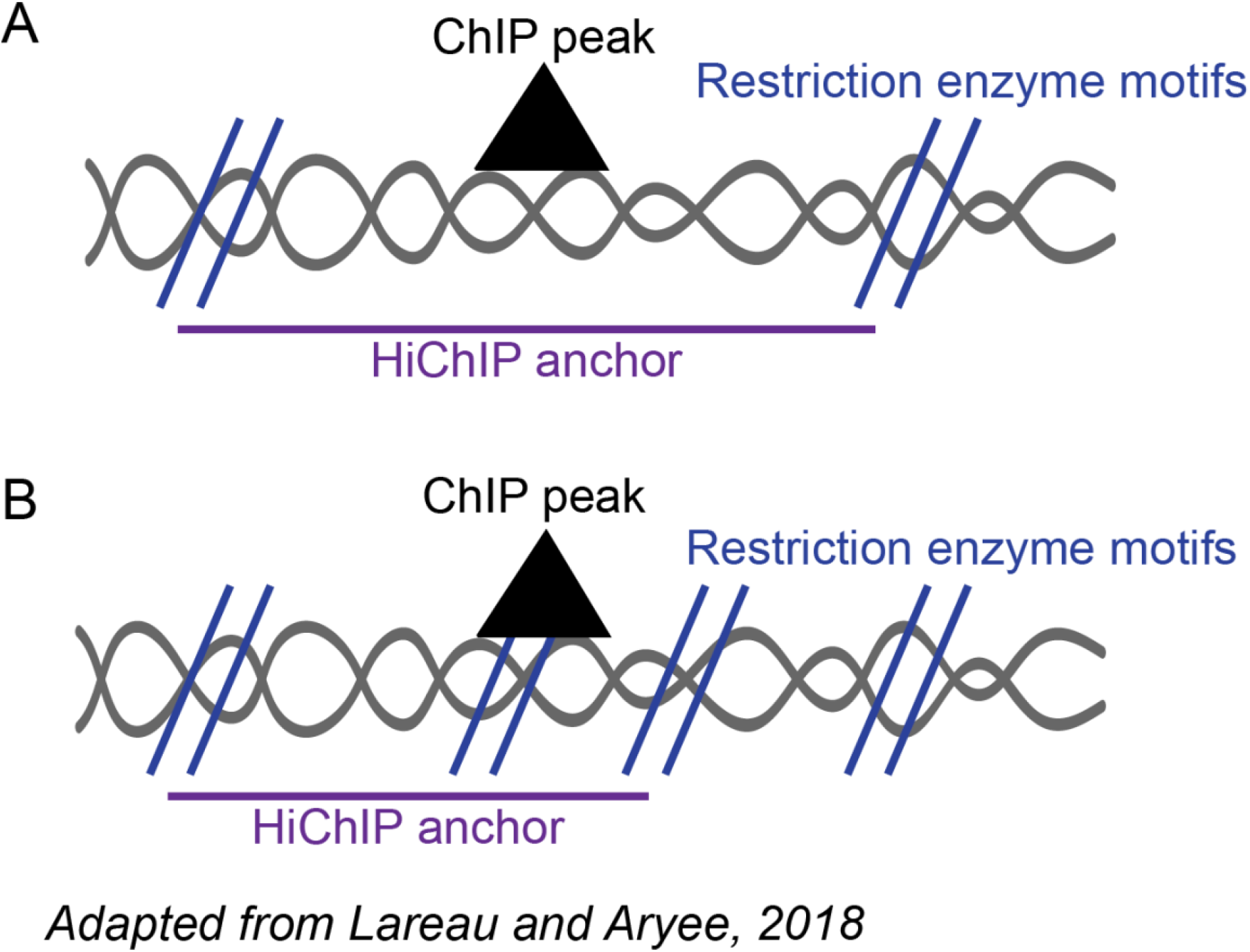
Identification of HiChIP anchors in hichipper. Related to Figure 4. (A) Diagram of restriction fragment-aware anchor calling when a ChIP peak is within a single restriction fragment. (B) Diagram of restriction fragment-aware anchor calling when a ChIP peak spans more than one restriction fragment. The anchor is then extended to include the length of overlapping fragments.

**Supplemental Figure 6.**
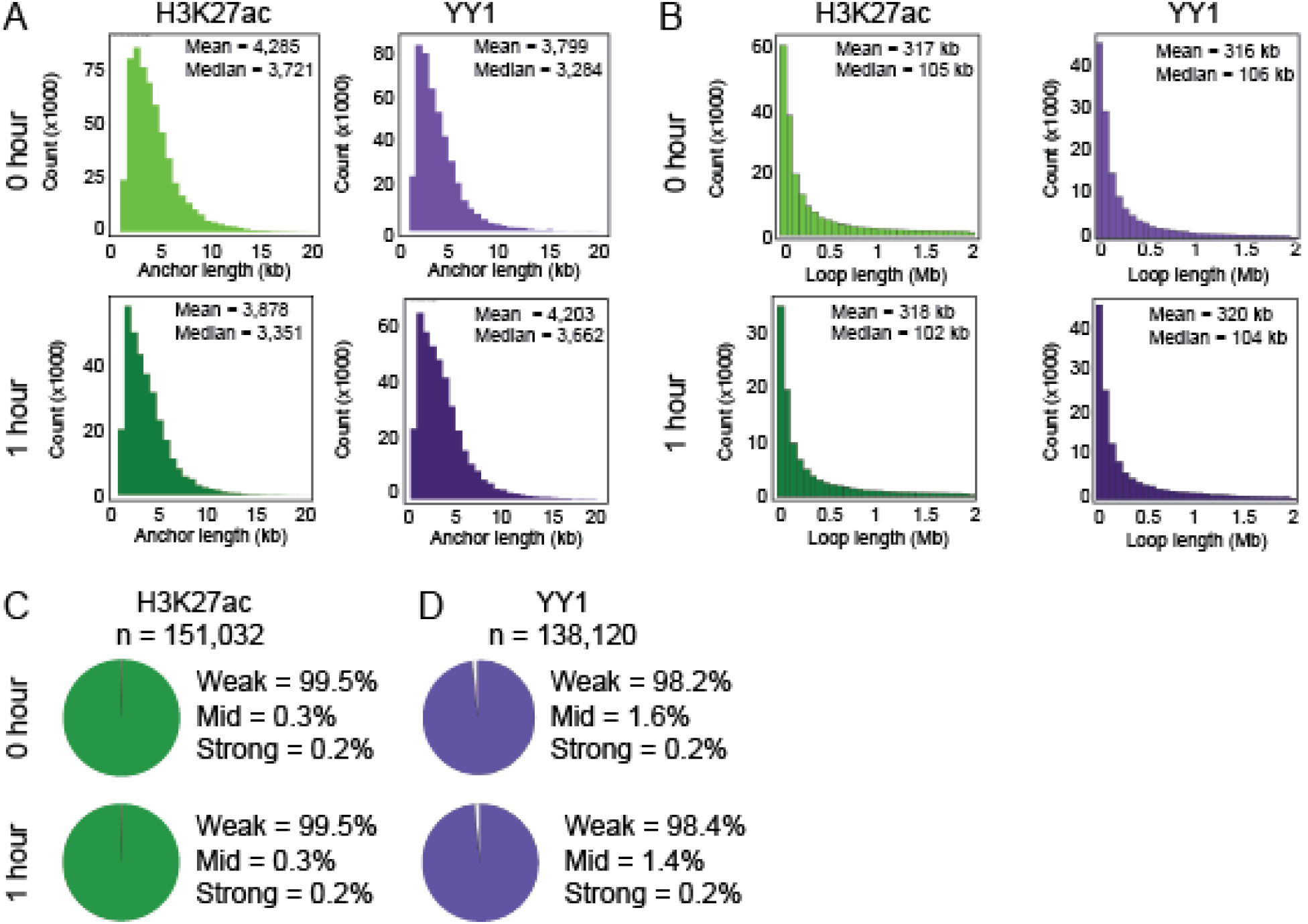
Characterization of chromatin loops mediated by H3K27ac and YY1. Related to Figure 4. (A) Histogram showing the size distribution of anchors for HiChIP loops in H3K27ac and YY1 libraries pre and post Epo stimulation. (B) Histogram showing the size distribution of HiChIP loops in H3K27ac and YY1 libraries pre and post Epo stimulation. (C) Fraction of weak (score < 5), mid (score between 5 and 10), and strong (score > 10) H3K27ac chromatin interactions. (D) Fraction of weak (score < 5), mid (score between 5 and 10), and strong (score > 10) YY1 chromatin interactions.

**Supplemental Figure 7.**
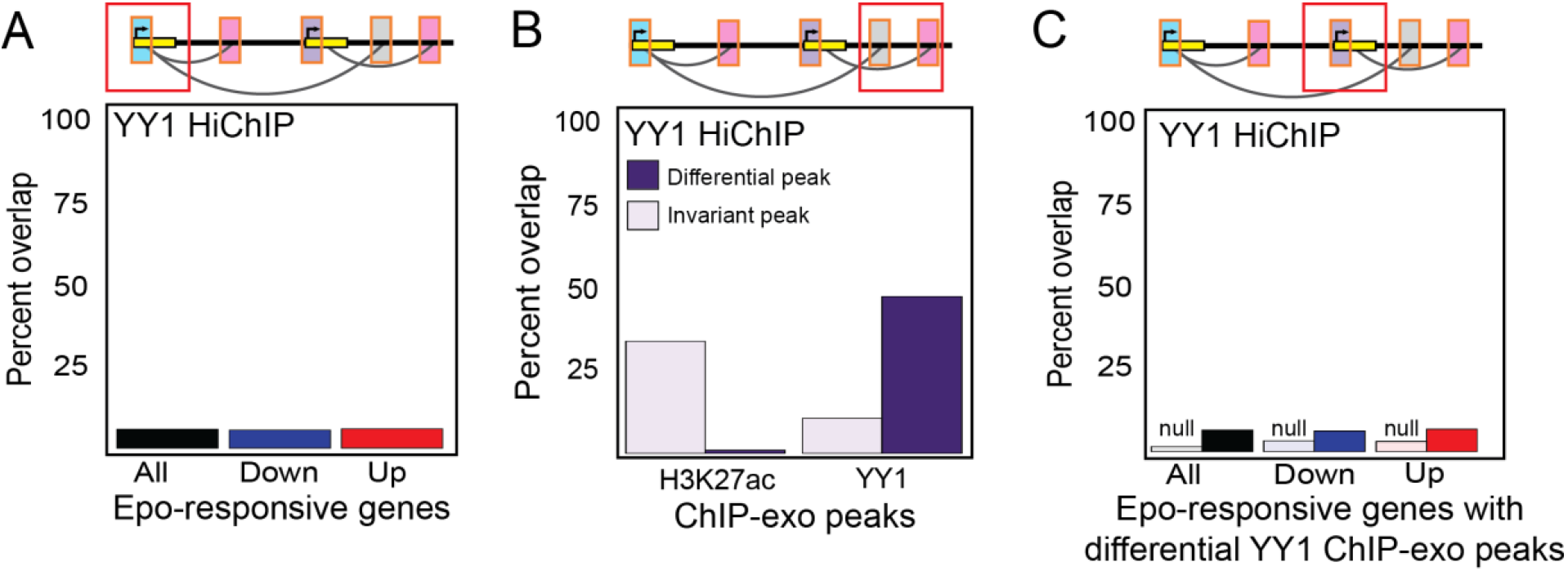
Epo regulates transcription in a pre-established chromatin conformation. Related to Figure 4. (A) Proportion of interactions with promoters of Epo-responsive genes within YY1 HiChIP anchor regions. (B) Proportion of interactions with differential H3K27ac or YY1 ChIP-exo peaks within anchor regions of YY1 HiChIP. Dark bars represent differential ChIP-exo peaks TSS and light bars represent invariant ChIP-exo peaks. (C) Proportion of interactions with differential YY1 ChIP-exo peaks at promoters of Epo-responsive genes within anchor regions of YY1 HiChIP. Dark bars represent Epo-responsive genes and light bars represent non-responsive genes.

## Acknowledgements

A.A.P. was supported by the Vanderbilt Molecular Endocrinology Training Program grant 5T32 DK07563. We would like to thank the Vanderbilt Technologies for Advanced Genomics (VANTAGE) Core for technical support with Illumina sequencing. Special thanks also go to Nicholas Servant, Maxwell Mumbach, and Caleb Lareau for experimental and computational assistance.

## Author Contributions

Conceptualization, A.A.P. and B.J.V.; Methodology, A.A.P. and B.J.V.; Formal Analysis, A.A.P.; Investigation, A.A.P.; Resources, A.A.P.; Writing – Original Draft, A.A.P.; Writing – Review and Editing, A.A.P., J.D.B., and B.J.V.; Supervision, B.J.V. and J.D.B.

