## Supplementary figures and images for "Erythropoietin regulates transcription and YY1 dynamics in a pre-established chromatin architecture"

### Supplemental Figure 1

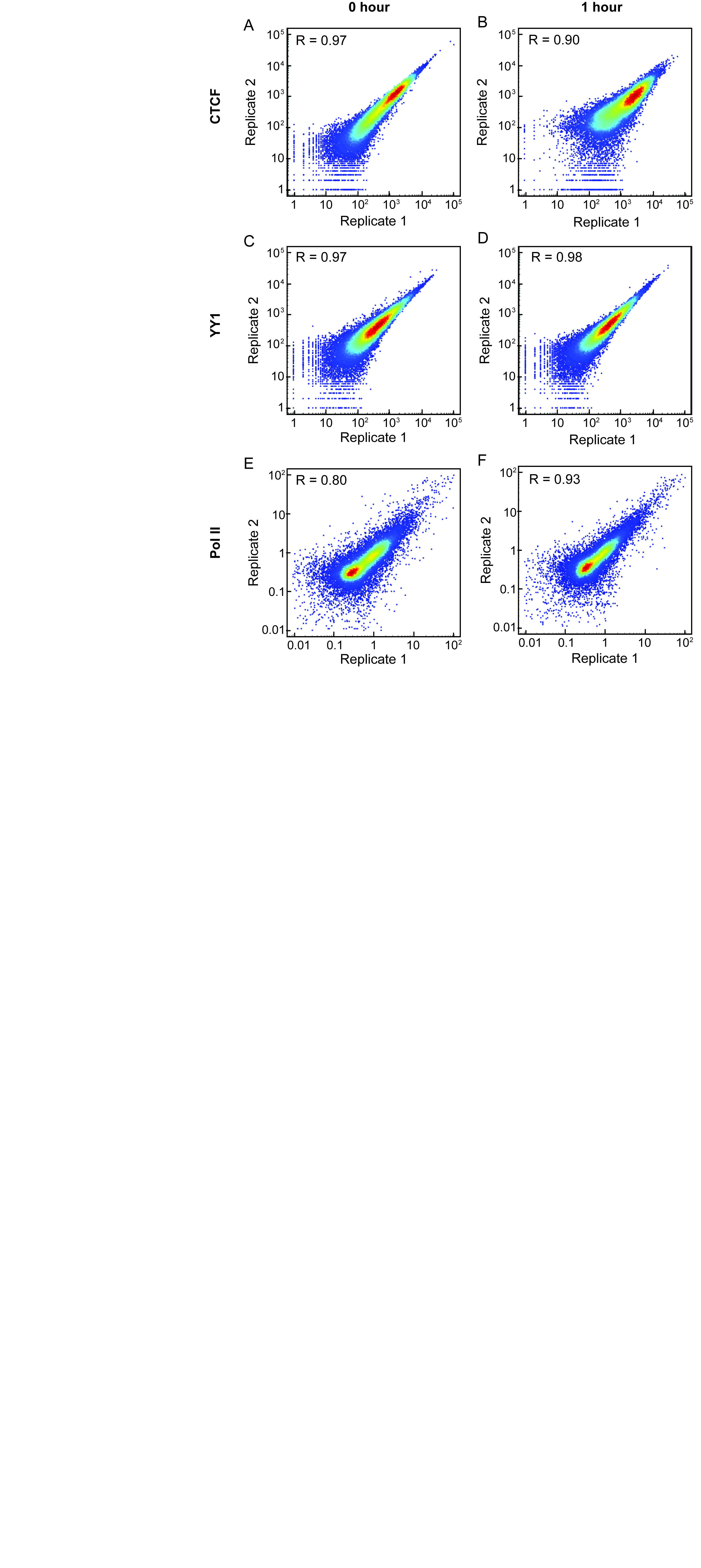

### Supplemental Figure 3

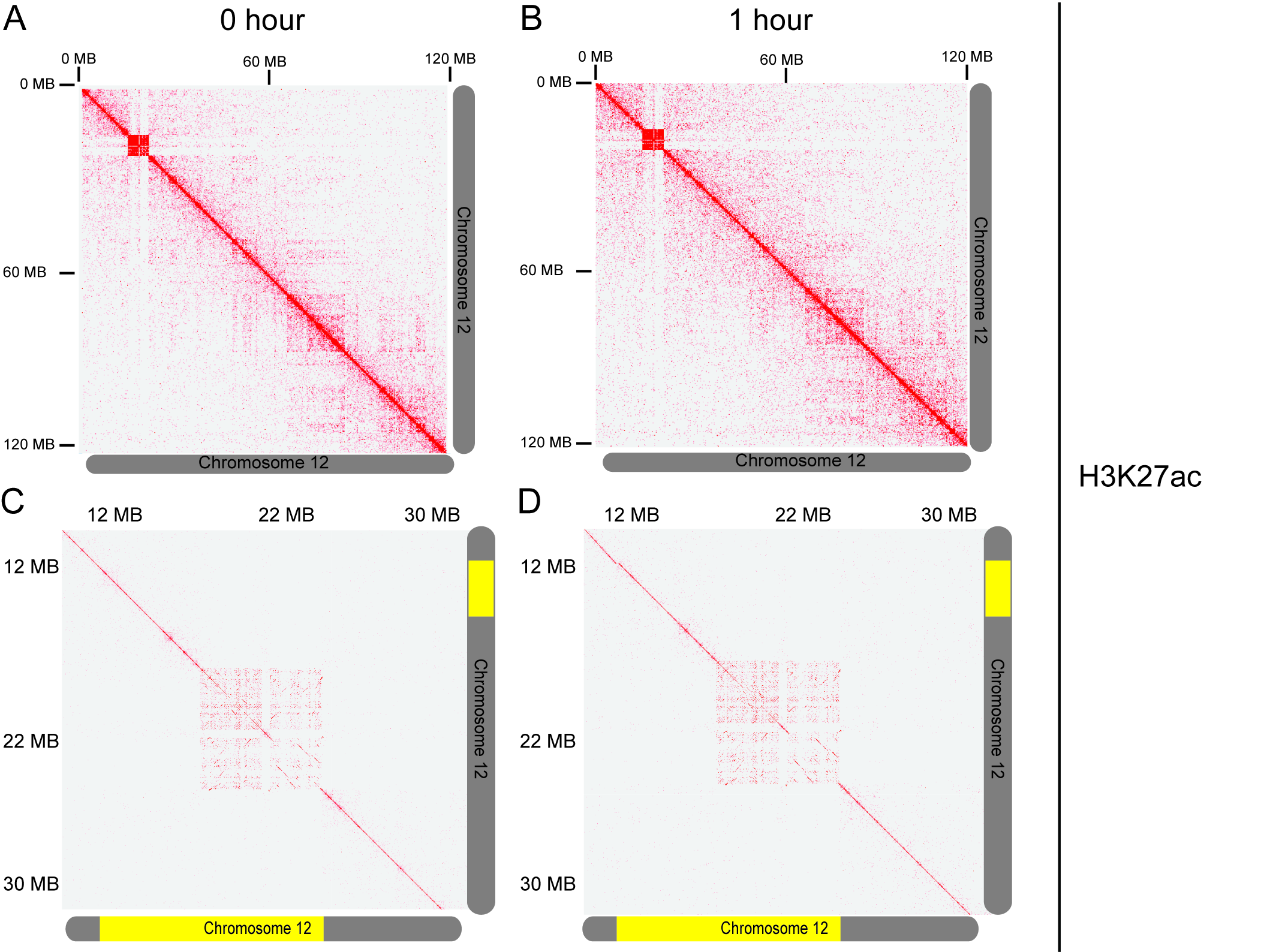

### Supplemental Figure 4

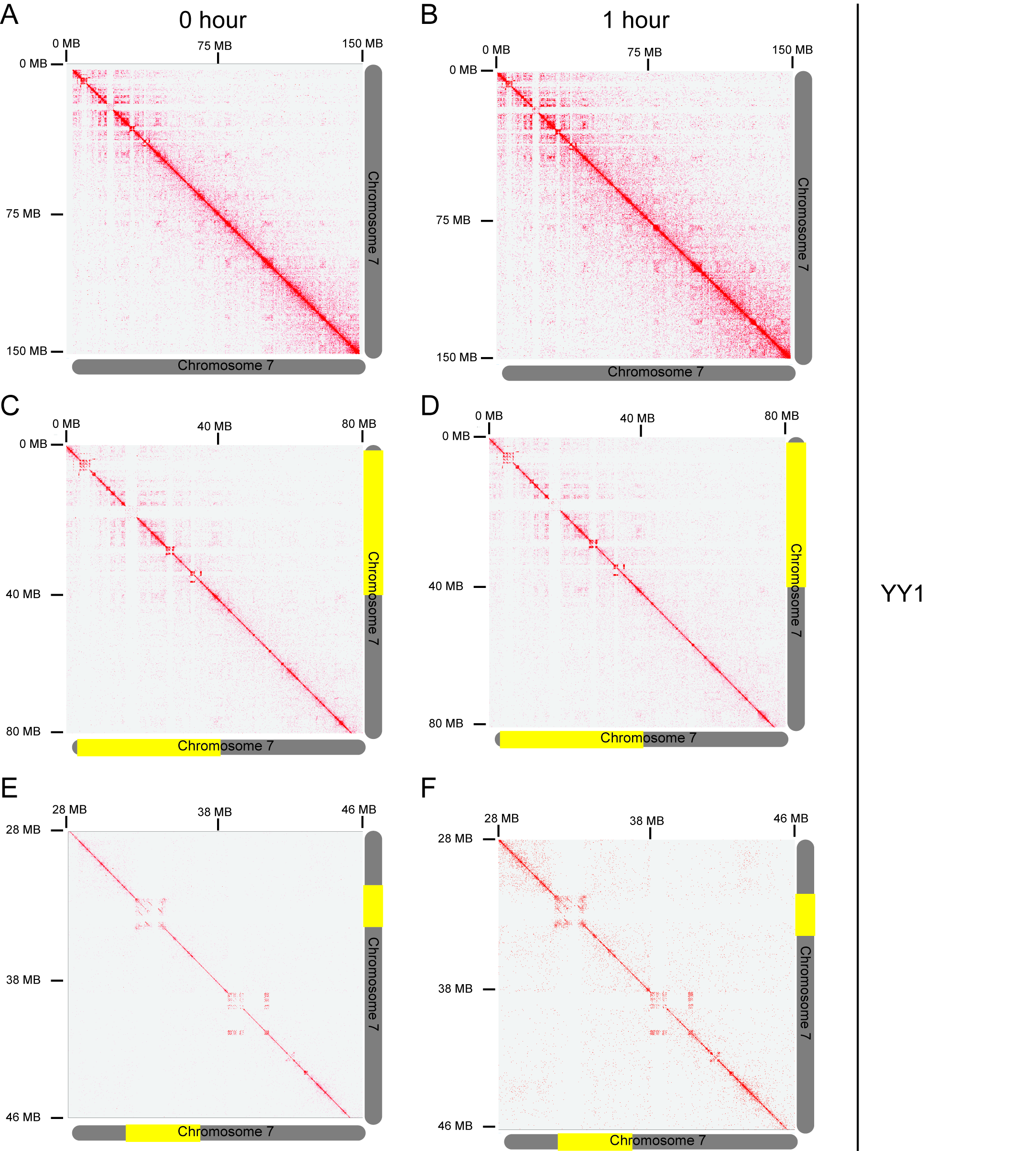

### Supplemental Figure 5

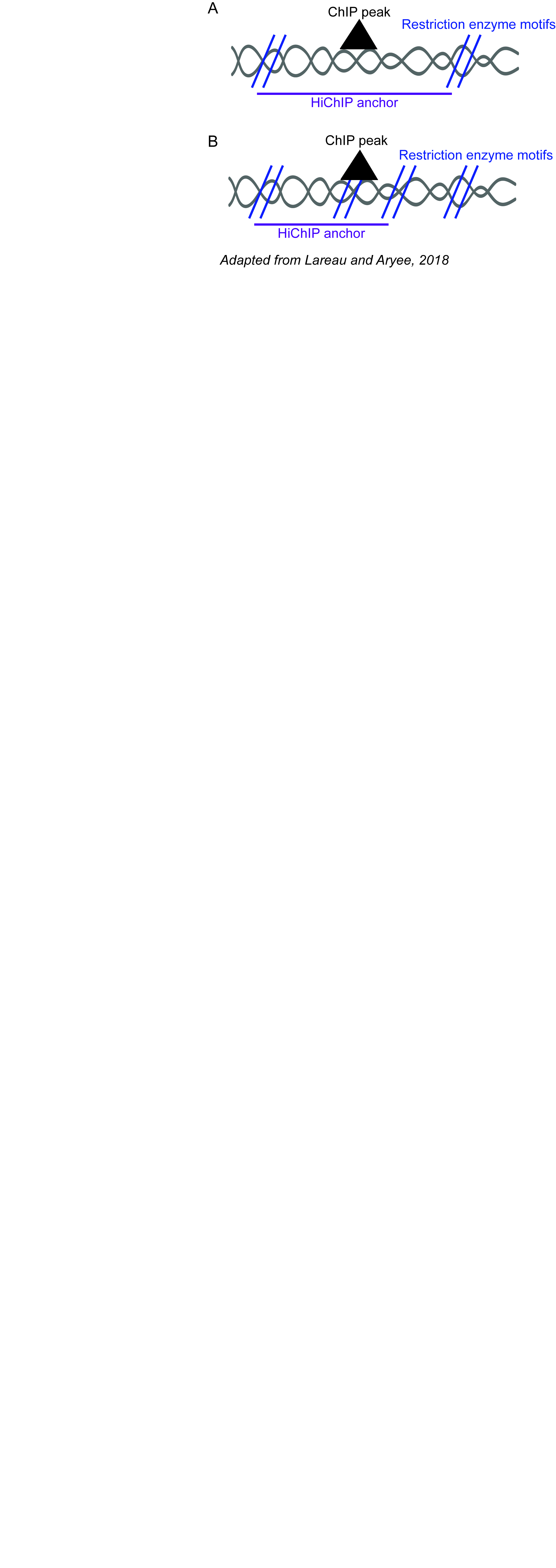

### Supplemental Figure 6

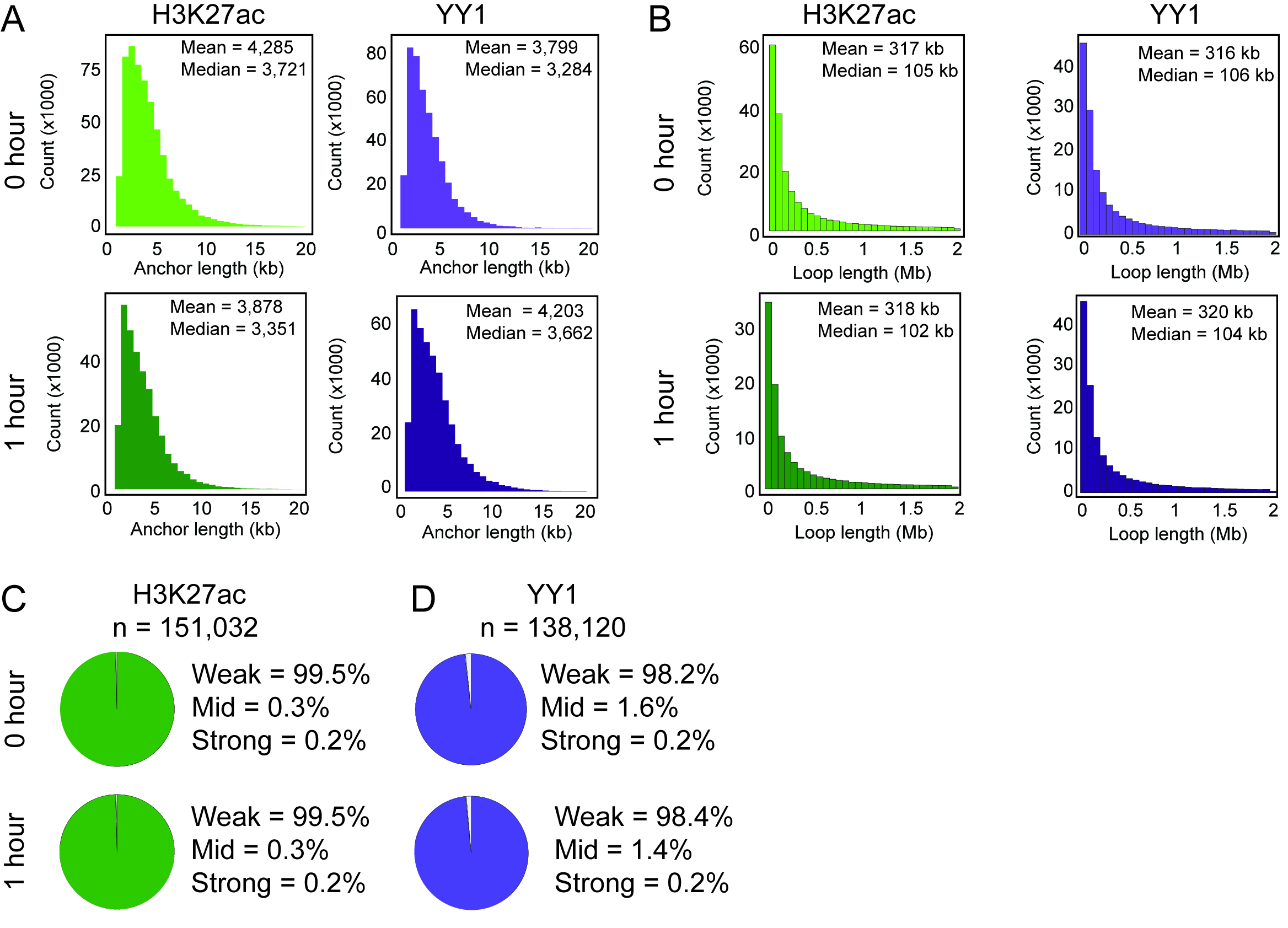
